# Cell polarity of neural crest-derived mesenchymal cells controls craniofacial development

**DOI:** 10.1101/2025.03.25.645191

**Authors:** Andrea Burianova, Kleopatra Kythraiotou, Lucie Sazavska, Ondrej Machon

## Abstract

Cell polarity is essential for tissue organization during development. While the molecular mechanisms governing epithelial cell polarity are well understood, the polarity of mesenchymal cells remains under debate. Using a mouse model with *Wnt5a* deletion in cranial neural crest cells, we analyzed polarized intracellular components that regulate mesenchymal cell organization, proliferation, and migration prior to mesenchymal condensation in the craniofacial region. Loss of *Wnt5a* disrupted the polarized localization of primary cilia and the Golgi apparatus along the proximal-distal axis. This intracellular disorganization impaired the orientation of the cell division plane and was accompanied by a reduced proliferation rate. As a result, several distal facial structures, including the frontonasal cartilage, premaxilla, and distal mandible, were significantly reduced in *Wnt5a* conditional mutants.

## INTRODUCTION

The craniofacial region is a defining feature of vertebrates, consisting of a complex arrangement of skeletal, muscular, and connective tissues that shape the face and head. Cranial neural crest cells (NCCs) play a pivotal role in forming craniofacial cartilage, bone, and connective tissue (Minoux and Rijli, 2010). The Wnt signaling pathway is a key regulator of NCC development. The canonical (β-catenin-dependent) signaling controling NCC induction and differentiation (García-Castro et al., 2002; Hari et al., 2002; Hari et al., 2012; Masek et al., 2016), while the noncanonical (β-catenin-independent) pathway, largely activated by WNT5A, is involved in the elongation of the anterior-posterior and proximal-distal body axes (Yamaguchi et al., 1999). The noncanonical WNT5A signaling regulates the planar cell polarity (PCP) pathway, which is essential for cell migration, cell polarity, and tissue organization in multiple embryonic regions including the craniofacial area. WNT5A signaling specifically triggers phosphorylation of the co-receptor ROR2, intracellular DVL2 and VANGL2 to establish the cell polarity (Gao et al., 2011; Ho et al., 2012; Konopelski Snavely et al., 2023). *Wnt5a* mutant embryos exhibit marked craniofacial abnormalities, including significant shortening of the snout and mandible, cleft palate, and reduced tongue length (He et al., 2008; Liu et al., 2012; Yamaguchi et al., 1999). In humans, mutations in the WNT5A, DVL, ROR2, or FZD2 genes cause craniofacial abnormalities characteristic of Robinow syndrome (Konopelski Snavely et al., 2023). Interestingly, certain dog breeds with shortened snouts also carry mutations in DVL2 (Mansour et al., 2018).

Within the forming cartilage, WNT5A signaling affects oriented cell division both in limb growth plates and in sheet-like cartilage structures in the head (Kaucka et al., 2017; Li et al., 2017). However, face formation and its shape seem to be already determined before cartilage formation. Lineage tracing experiments showed that cranial neural crest cells populate facial area in cell clusters representing clones of mesenchymal cells. Again, noncanonical WNT5A signaling participates in determining oriented cell division of mesenchymal cells and in orchestrating local directional cell movements to define face shaping. (Kaucka et al., 2016b). Although the increasing evidence supports the involvement of WNT5A/PCP pathway in shaping the craniofacial region, very little is known about molecular determinants of the cell polarity of NCC-derived mesenchyme. Studies suggest that these mesenchymal cells utilize a specific set of PCP components that may differ from other cellular systems. For instance, VANGL1/2 proteins are essential for the neural tube closure but they appear dispensable for NCC migration in mouse (Pryor et al., 2014).

PCP pathway is known to affect front-rear polarization to direct their movement by rearranging the cytoskeleton, junctional remodeling and by modulating extracellular matrix (ECM) interacting proteins (Koca et al., 2022; Ladoux et al., 2016). It is largely unknown what molecular mechanisms are specifically involved in directing migration of NCC-derived murine mesenchymal cells. Studies in *Xenopus* and *Danio* suggest that Rho and Rac small GTPases mediate non-canonical Wnt signaling to arrange cytoskeletal components during front-rear cell polarization (Matthews et al., 2008). Although the role of PCP-linked GTPases is described well during epithelial-mesenchymal transition and NCC emigration from the neural tube (Mayor and Theveneau, 2014), the data on PCP function in craniofacial mesenchymal cells are limited. NCC-specific deletion of Rac1 using Wnt1-Cre driver resulted in severe mid-facial cleft (Thomas et al., 2010), while activation of dominant-negative form of *Rock* by the same driver led to frontonasal and mandible hypoplasia (Phillips et al., 2012). Moreover, directional migration of enteric NCC depends on accumulation of αE-Catenin, β-Catenin and Cadherins in lamellipodia in the front end which is accompanied by polarized distribution of RhoA and cytoskeletal myosin-IIB (Vassilev et al., 2017). These findings raise several important questions. How is the polarity of NCC-derived craniofacial mesenchymal cells determined? What cytoskeletal and cell surface components are involved in front-rear polarization that control interaction with ECM and neighboring cells and affect directional cell movement? How do these processes ultimately determine face shaping? The cellular mechanisms governing the early steps preceding cartilage growth are not fully understood, such as migration and spatial organization of chondrogenic neural crest cells into the craniofacial region, distribution of new mesenchymal condensations and their integration into the growing cartilage, or initiation of intramembranous ossification. It is not clear what target genes are involved in these developmental processes, what cellular interactions determine the ultimate shape of the face, more specifically the shape of bones and cartilage. Non-canonical WNT pathway regulates both transcription of target genes as well as the cytoskeletal architecture. In the context of craniofacial development, it is still unclear what branch of WNT/PCP signaling controls transcription and what branch directly modulates cytoskeletal reorganization and cell polarity. The cell polarity is well described in epithelial structures, but the polarity of mesenchymal cells is not sufficiently characterized. It remains to be described what cellular components determine mesenchymal cell polarity, how these factors differ from epithelial cells and what mechanisms regulate mesenchymal cell organization into functional tissues. So far, Wnt/PCP pathway was reported to be implicated in contact-mediated inhibition of locomotion of neural crest cells (Carmona-Fontaine et al., 2008; Davey and Moens, 2017) , and in chondrocyte polarity (Wan and Szabo-Rogers, 2021). Nonetheless, critical molecular components of the spatial arrangement of craniofacial mesenchymal cells are largely unknown.

Using a conditional knockout model targeting *Wnt5a* in NCCs, we analysed key cellular processes controlling morphogenesis in the craniofacial region. We identified determinants of cell polarity in mesenchymal cells in the developing frontonasal process (FNP) and the first pharyngeal arch (PA1) that are under the control of non-canonical Wnt/PCP pathway. Thus, polarized localization of primary cilia, centrosomes and Golgi apparatuses together, together with the oriented cell divisions and proliferation rate, are the key factors forming the shape of craniofacial region.

## RESULTS

### Severe craniofacial defects in Wnt1-Cre2; *Wnt5a fl/fl* embryos result from reduced cell proliferation and initial abnormal shaping of the frontonasal process and the pharyngeal arch

To study the role of WNT5A signaling specifically in the neural crest derived structures in the craniofacial region, we employed Wnt1-Cre2 mouse (Lewis et al., 2013). It is important to note that all mesenchymal cells in FNP and PA1 are targeted by Wnt1-Cre2 whereas all epithelial structures are not (Hudacova et al., 2024; Lewis et al., 2013; Schock et al., 2017).

High-resolution chain reaction fluorescent *in situ* hybridization (HCR FISH) revealed that *Wnt5a* was strongly expressed in FNP at E10.5 while PA1 showed weaker expression at this stage (Fig. 1A-A’, arrows). At E11.5, *Wnt5a* expression appeared strong in distal parts of both FNP and PA1 (Fig. 2B). We crossed Wnt1-Cre2 mouse strain with Wnt5a ^fl/fl^ strain to obtain Wnt1-Cre2; *Wnt5a fl/fl* (*Wnt5a* cKO). Wnt5a HCR FISH on frozen sagittal sections together with gross morphology inspection revealed that both FNP and PA1 were smaller and remarkably shorter along the proximal-distal (P-D) axis in mutants already at E11.5 (Fig. 1B-B’ arrows and small insets up, right). In contrast to systemic *Wnt5a* KO embryos (Yamaguchi et al., 1999), in which *Wnt5a* was deleted in the whole body, the trunk region and limbs appeared normal (not shown). Shortening and widening of FNP was even more conspicuous as development progressed.

**Fig. 1.**
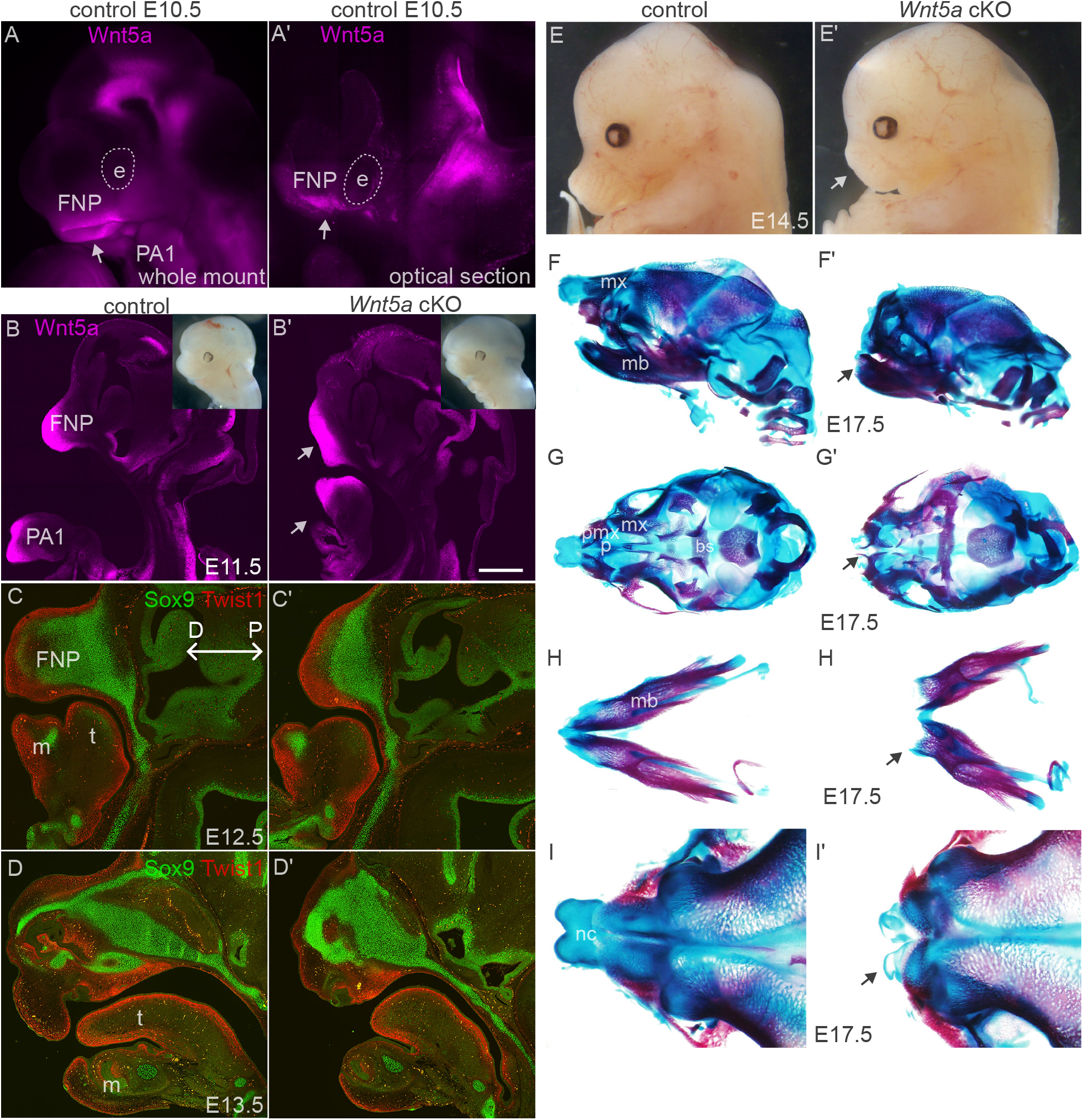
Drastic malformations of distal bones and cartilages in *Wnt5a* cKO. (A-A’) Whole-mount HCR FISH of *Wnt5a* transcripts at E10.5 embryonic heads. (A’) shows its optical section (one z-stack). (B-B’) *Wnt5a* mRNA HCR FISH on frozen 100-μm thick sagittal sections of E11.5 embryonic heads of control and *Wnt5a* cKO. Small insets (right, up) show shortening of distal craniofacial region in the bright-field light. (C-D’) Immunostaining of paraffin sections of control (C, D) and Wnt5a cKO (C’, D’) at E12.5 and E13.5, demonstrating the expression patterns of these cell populations: Sox9 antibody (green) labels prechondrogenic condensations and chondrocytes. Twist1 (red) antibody marks the localization of Twist1^+^ NC-derived mesenchymal subpopulation. Proximal-Distal (P-D) axis is illustrated in (C). (E-E’) Side views of E14.5 control (F) and *Wnt5a* cKO (F’) embryos in the bright-filed light. Note a shortened frontonasal region (white arrowhead). (F-I’) Alcian Blue and Alizarin Red staining at E17.5, visualizing the cartilage (blue) and bone (purple). (F-F’) Lateral views of control (F) and *Wnt5a* cKO (F’) skulls. Note a missing distal mandible (black arrowhead). (G-G’) Ventral views of control and *Wnt5a* cKO crania. Note a reduced premaxilla and bifid, as well as a shortened nasal cartilage in the mutant (black arrow in G’). (H-H’) Dorsal views of control and *Wnt5a* cKO mandibles. Note a size reduction and widening in the the mutant (black arrowhead in H’). (I-I’) Dorsal views of control and *Wnt5a* cKO nasal regions. Note a massive reduction of the nasal bone and a shortened bifid nasal cartilage in the mutant (black arrow in I’). bo; basioccipital; bs, basisphenoid; cn, cartilage primordium of the nasal bone; f, frontal; FNP, frontonasal process; m, mandible; mx, maxilla; na, nasal; nc, nasal cartilage; np, nasal process; p, parietal; PA1, the first pharyngeal arch; pmx, premaxilla; ps, presphenoid; t, tongue. Scalebars: 200 μm

**Fig. 2.**
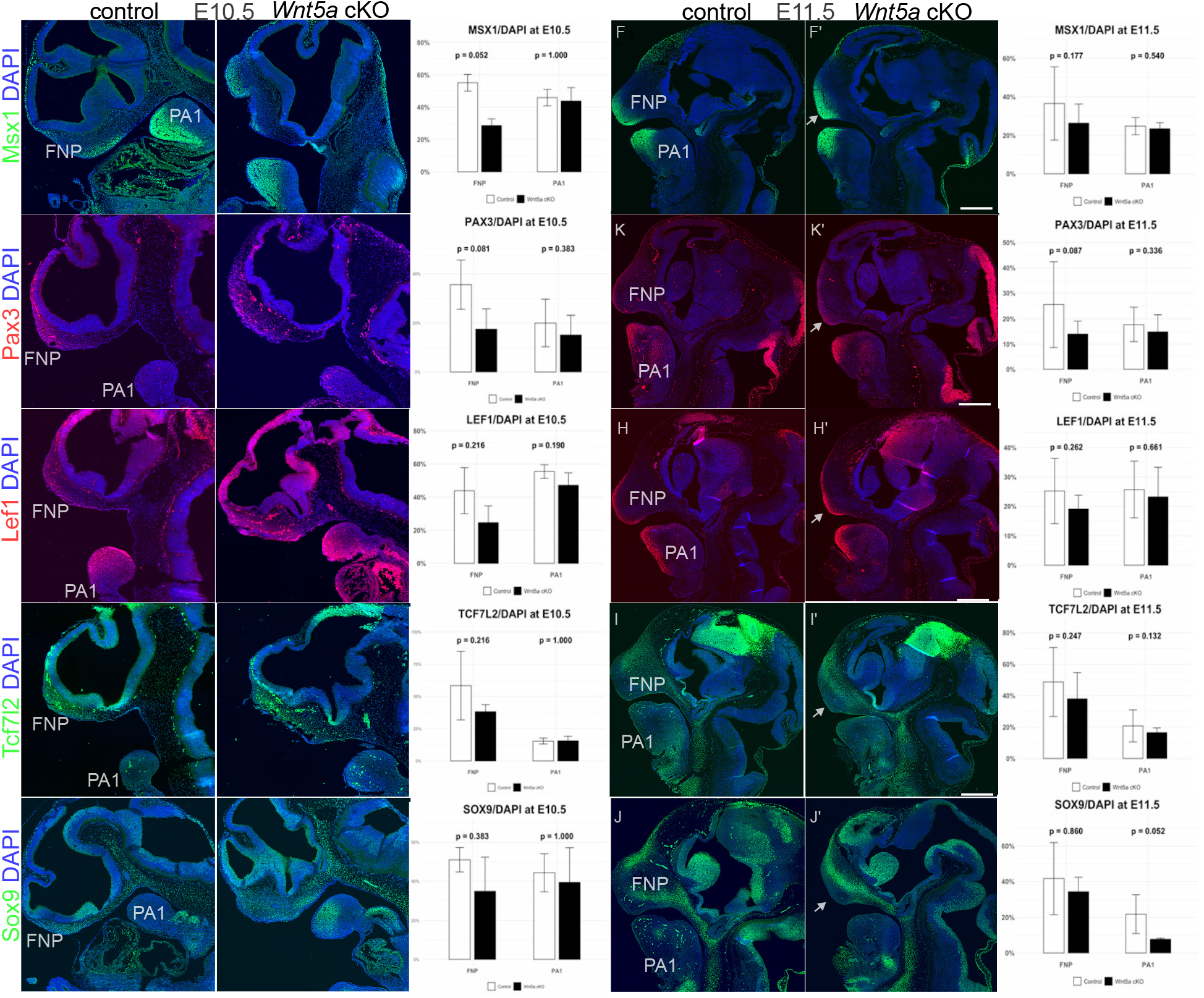
Severe shortening of the distal craniofacial region in *Wnt5a* cKO. (A-A’’) Fluorescent immunohistochemistry of sagittal paraffin sections of control and *Wnt5a* cKO heads using Msx1 antibody, with accompanying quantification graphs (A’’). (B-B’’) Pax3 staining. (C-C’’) Lef1 staining. (D-D’’) Tcf7l2 staining. (E-E’’) Sox9 staining. (F-J’) Immunostaining of paraffin sagittal sections of control and *Wnt5a* cKO heads at E11.5 with the same set of antibodies. A’’-J’’ quantification of cell markers relative to DAPI+ nuclei for each cell population at E10.5 and E11.5. FNP, frontonasal process; PA1, the first pharyngeal arch. Scalebars: 200 μm

Analysis of sagittal paraffin sections of embryos at E12.5-13.5 confirmed severe reduction of the distal FNP and PA1 (Fig. 1C-D’). Interestingly, superficial mesenchyme labelled with Twist1 antibody and prechondrogenic mesenchyme labelled with Sox9 antibody were specified properly despite severe morphological defects. Thus, FNP and PA1 shortening already at E11.5 led to failure of facial extension along P-D axis at E14.5 and later (Fig. 1E-E’). Analysis of bones and cartilages at E17.5 by Alcian and Allizarin staining documented that distal craniofacial structures were strongly reduced. The nasal cartilage and premaxilla were almost absent, and a distal part of the mandible was very short (Fig. 1F-I’). Moreover, we also noticed defects in more proximal structures, e.g. the cleft palate and malformed basisphenoid bone (Fig. 1G’). In summary, dramatically malformed or almost missing distal craniofacial areas in Wnt5a cKO document indispensable function of *Wnt5a* distally which also corresponds to its spatially restricted expression in these regions.

Next, we analysed expression of multiple cellular markers to examine cell specification along the P-D axis using immunostaining at the onset of phenotypic changes at E10.5 and E11.5 (Fig. 2). To quantify the number of cells in each population, we counted cell numbers relative to DAPI-labelled nuclei in FNP and PA1 compartments. Expression of all measured genes (*Msx1, Pax3, Lef1, Tcf7l2, Sox9*) was reduced in *Wnt5a* cKO to almost one half in FNP at E10.5 and in E11.5. Interestingly, downregulation of these markers in PA1 became measurable at E11.5 but not earlier (see quantification panels to the right in Fig. 2). This may correspond to a weaker expression of *Wnt5a* at PA1 compared to FNP when analysed at E10.5 (see Fig. 1A’). At E11.5, the most distal regions of FNP and PA1 labelled with MSX1, PAX3 and LEF1 antibodies appeared shorter and wider reflecting the morphological changes in *Wnt5a* cKO (Fig. 2F-H’, arrows). Interestingly, all these cell markers were maintained suggesting that cell fate was not much affected in distal FNP and PA1 in the mutants, but these regions changed size and shape. TCFL2 protein labels the prechondrogenic mesenchyme at E11.5 in the proximal FNP because this area also strongly expresses *Sox9* (Fig. 2I-J). The size of this proximal part was also reduced in *Wnt5a* cKO. The number of TCF7Ll2+ cells decreased from to 58% to 38% while the number of Sox9+ cells decreased from 51% to 35% in the mutant FNP at E11.5 (Fig. 2I’’ and 2J’’). The overall decreased number of cells labelled with specific markers indicated that cell proliferation was affected in *Wnt5a* cKO mutants. To analyse proliferation rate, we injected EdU into pregnant mice 2 hours prior to harvesting and calculated the number of cells in S-phase relative to DAPI. In FNP of controls at E10.5, the number of EdU+ cells was calculated on average 55% of all DAPI cells, whereas the ratio decreased to 46% in *Wnt5a* cKO mutants. In PA1 at E10.5, the EdU ratio decreased from 51% to 45% in mutants (Fig. 3A-A’’). At E11.5, the proliferation rate in FNP was calculated to decrease from 36% in controls to 27% in mutants (Fig. 3B-B’’).

**Fig. 3.**
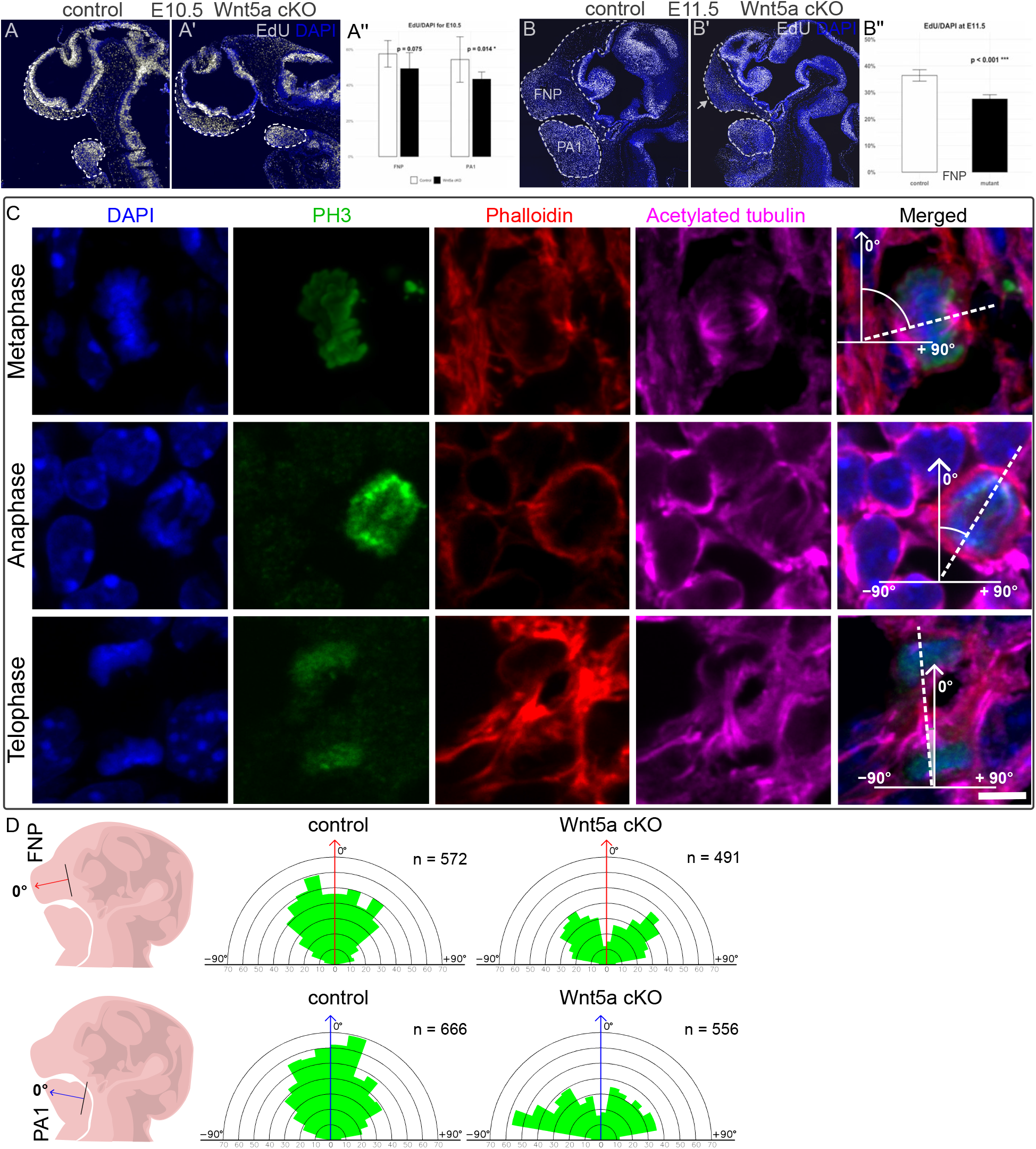
Reduced proliferation rate and disarranged orientation of cell division plane in *Wnt5a* cKO. (A-A’’) Short-pulse (2-hour) EdU incorporation in the embryonic heads at E10.5, sagittal sections with a quantification graph (A’’). (B-B’’) Short-pulse EdU incorporation at E11.5 with a quantification graph (B’’). (C) Immunostaining of mitotic cells with PH3, Phalloidin and Acetylated-tubulin antibodies illustrating measurements of division plane orientations. The angle 0 degrees represents the line parallel to the arbitrary P-D axis in FNP and PA1 schematically shown in (D, left). (D) Rose diagrams showing quantifications of the division plane orientation relative to P-D axis in FNP and PA1. n, cell counts for each condition. Scalebar: 5 μm

### Cell polarity and oriented cell division of mesenchymal cells derived from cranial NCC

Previous studies suggested that Wnt/PCP pathway affects orientation of the cell division plane in the craniofacial region potentially explaining shorter but wider snout of *Wnt5a* KO embryos (Kaucka et al., 2016a). Similarly, disrupting Wnt/PCP pathway in the already formed chick limb cartilage using retroviral vectors resulted in loss of oriented cell division of chondrocytes and impaired directional growth (Li et al., 2017). To investigate the division plane orientation in FNP and PA1, we used a combination of three immunohistochemical markers at E11.5: PH3 to label mitotic cells, Alexa-coupled phalloidin to visualize actin filaments and cell shape, and acetylated tubulin to detect mitotic spindles (Fig. 3C). P-D axes in FNP and PA1 were arbitrarily defined as depicted in the schemes in Fig. 3D, with the angle set to 0 degrees. By measuring more than 500 cells for each condition we found that in the control group, most division axes aligned with P-D reference line, deviating by a maximum of ±40^0^ both in FNP and PA1 (Fig. 3D). In contrast, in *Wnt5a* cKO embryos, this preferential orientation of division was lost, with a broader distribution of angles both in FNP and PA1 (Rose diagrams in Fig. 3D).

Interestingly, most mutant cells showed a mean deviation value around ±45^0^, with very few division axes near 0 degrees. This effect was more pronounced in PA1, where most dividing cells deviated by ±70-80^0^, suggesting a stronger impact of *Wnt5a* loss in the mandibular arch PA1. These results indicate that craniofacial mesenchymal cells have features of planar cell polarity at E11.5 which is reflected in the oriented cell division plane along the P-D axis.

### Polarized localization of primary cilia, Golgi apparatuses and centrosomes in craniofacial mesenchymal cells

In most epithelial cells, the apical positioning of primary cilia reflects the inherent polarity of epithelial cells, which organize distinct apical and basolateral domains separated by tight junctions. The apical location of the primary cilium positions enables sensing and responding to mechanical or chemical signals as well as cilium-dependent signaling pathways in epithelial tissues. We therefore tested the possibility whether mesenchymal cells also exhibit polarized localization of primary cilia. We visualized primary cilia using Arl13B antibody immunostaining (Fig. 4). All mesenchymal cells have primary cilia. We measured their angle towards the nucleus center (Fig. 4A’’) and relative to the arbitrary proximal-distal line schematically depicted in (Fig. 4B). Quantification of approx. 9000 cells showed that 62% and 65% of primary cilia are directed towards the distal pole of control FNP and PA1, respectively (Fig. 4B’). In *Wnt5a* cKO, however, this number decreased to 56% and 53% in FNP and PA1, respectively (Fig. 4B’’).

**Fig. 4.**
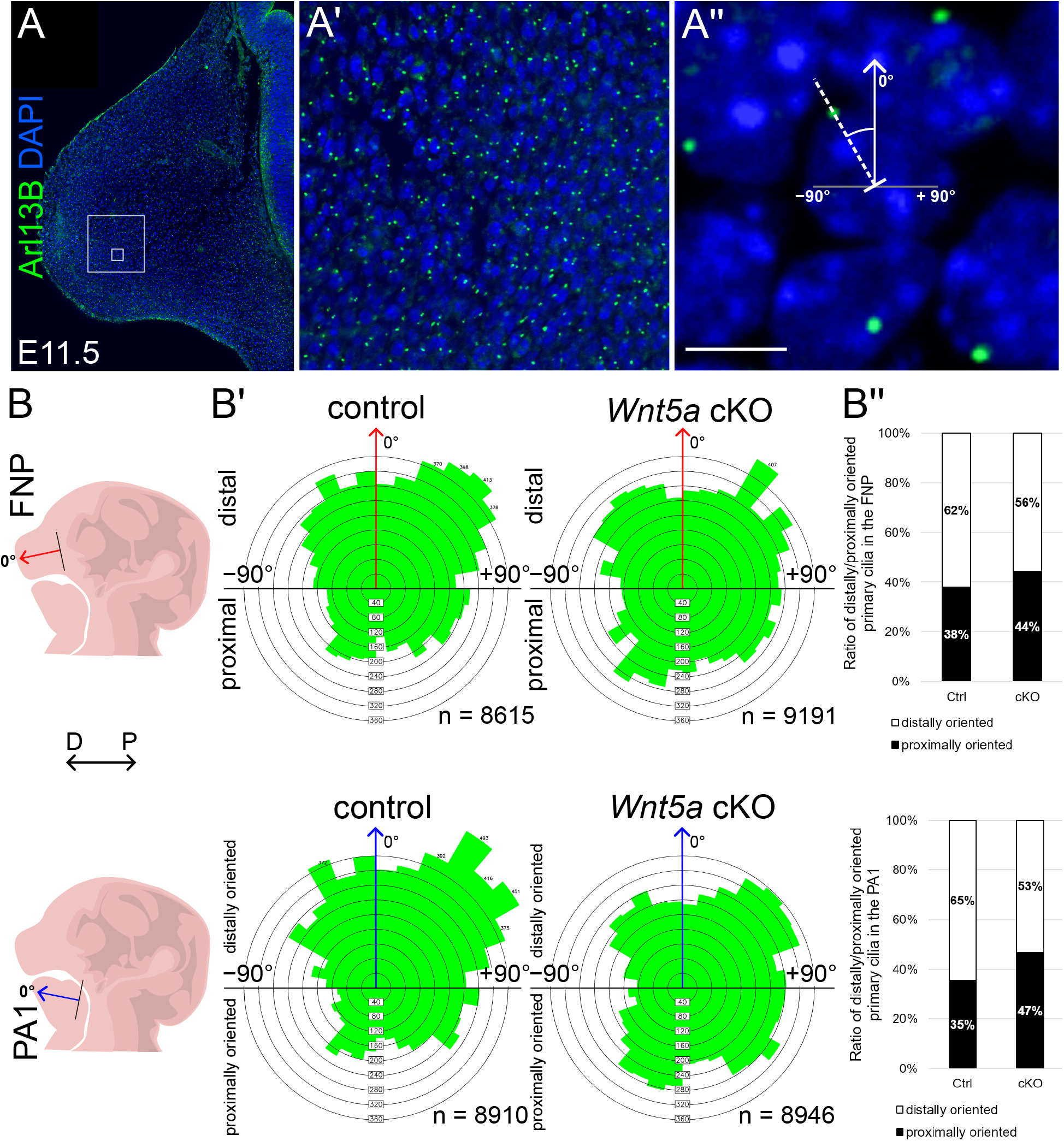
Directed orientation of primary cilia in craniofacial mesenchymal cells is impaired in *Wnt5a* cKO. (A-A’’) Immunostaining of primary cilia on sagittal sections at E11.5 using Arl13B antibody. (A’-A’’) show magnified areas depicted by rectangles in (A). The angle of cilia orientation was measured towards the centre of nucleus labelled with DAPI. The angle 0 degrees represents the line parallel to the arbitrary P-D axis in FNP and PA1 schematically shown in shown in (B, left). (B-B’) Rose diagrams summarizing quantifications of primary cilia angles relative to the nucleus centre in FNP and PA1. n, cell counts for each condition. (B’’) The ratio of distally and proximally oriented primary cilia in angles relative to the nucleus centre in FNP (top) and PA1 (bottom) in controls and mutants. Scalebar: 5 μm

Cell division orientation is affected by cytoskeleton organization, particularly by the position of centrosomes that serve as microtubule-organizing centres. Centrosomes also participate in forming a basal body of the primary cilium. To support our observation that primary cilia are preferentially oriented towards the distal end in mesenchymal cells, we analysed the position of Golgi apparatuses and centrosomes in FNP and PA1. Similarly to the primary cilia analysis, we measured the position of Golgi apparatus towards the arbitrary P-D axis and the nucleus centre using GM130 antibody (Fig. 5 A-A’’). The protein GM130 is a part of Golgi matrix. Quantification of the angles depicted in Fig. 5A’’ showed that most of Golgi apparatuses in control FNP were located distally within the angles -90^0^-30^0^. In contrast, most of *Wnt5a* mutant cells in FNP exhibited Golgi localization in the proximal sector within angels +90^0^-+135^0^ (Fig. 5B’). Next, we analysed centrosome positions using pericentrin antibody and measured again the angle towards the nucleus centre (Fig. 5D-D’’). However, the position of centrosomes did not show a tendency to specific orientation towards P-D axis (Fig. 5E-F’’).

**Fig. 5.**
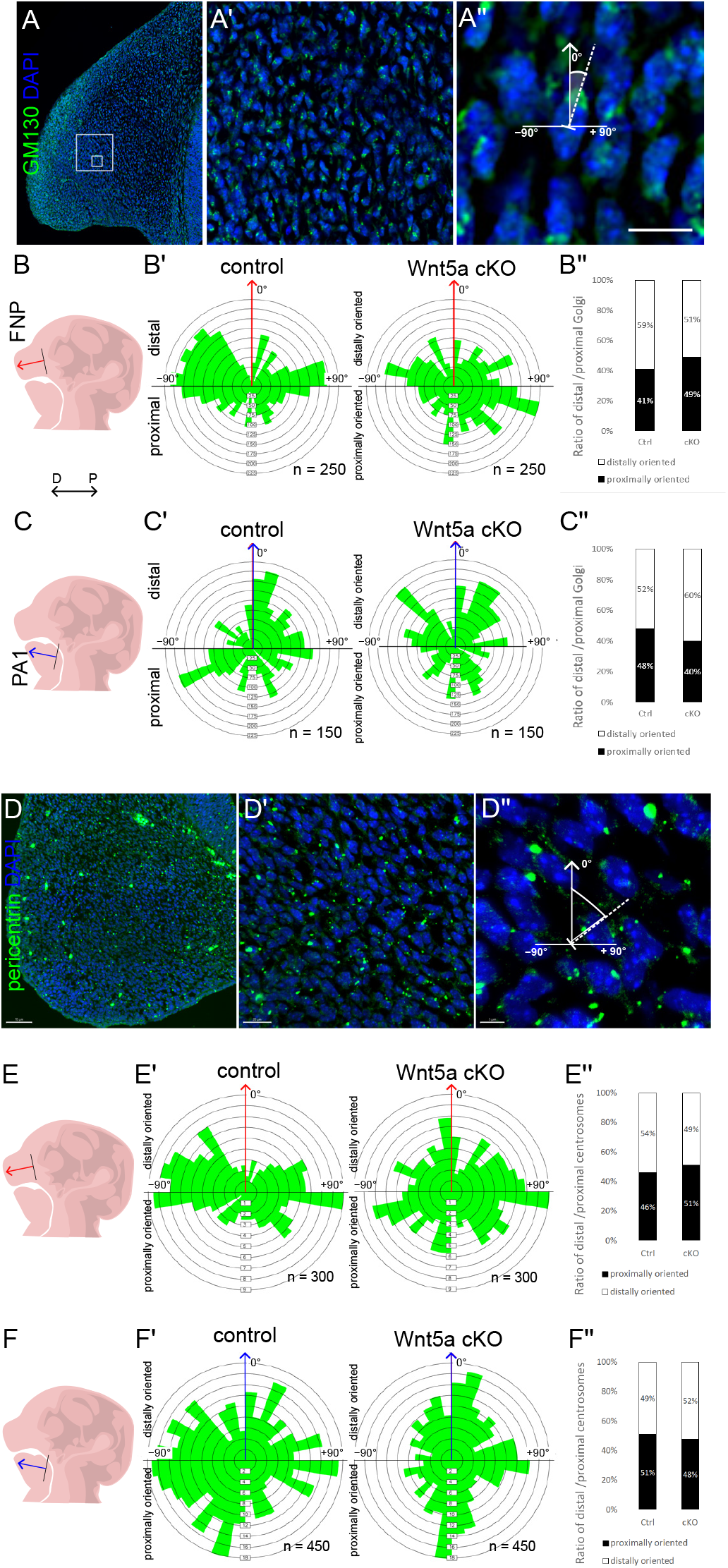
Disarranged orientation of Golgi apparatuses in *Wnt5a* cKO. (A-A’’) Immunostaining of Golgi apparatus using GM130 antibody. (A’-A’’) show magnified areas depicted by rectangles in (A). The angle of Golgi orientation was measured towards the centre of nucleus labelled with DAPI. The angle 0 degrees represents the line parallel to the arbitrary P-D axis schematically shown in (B-C, left). (B-C’) Rose diagrams summarizing quantifications of Golgi positions in angles relative to the nucleus centre in FNP and PA1 (C’). n, cell counts for each condition. (B’’) The ratio of distally and proximally oriented Golgi in FNP and PA1 (C’’). (D-D’’) Immunostaining of centrosomes using pericentrin antibody. (D’-D’’) show magnified areas depicted by rectangles in (D). The angle of centrosome orientation was measured towards the centre of nucleus labelled with DAPI. The angle 0 degrees represents the line parallel to the arbitrary P-D axis schematically shown in (E-F, left). (E-F’) Rose diagrams summarizing quantifications of centrosome positions in angles relative to the nucleus centre in FNP and PA1 (F’’). n, cell counts for each condition. (E’’) The ratio of distally and proximally oriented centrosome in FNP and PA1 (F’’).

## DISCUSSION

Wnt5a signaling has been described to affect craniofacial development a time ago (Yamaguchi et al., 1999). Systemic knock-out of *Wnt5a* in mice resulted in severe failure in the growth of limbs, extension of the anterior-posterior body axis as well as in drastic truncation of craniofacial structures. Although the consequences of *Wnt5a* deletion on mouse embryogenesis are well known, the molecular mechanisms underlying these develpmental defects are largely uknown. Here we focused on molecular characterization of mesenchymal cells derived from cranial NCC that primarily form the face and a part of the cranium. Our study provides compelling evidence that noncanonical Wnt5a signaling is essential for the establishment of mesenchymal cell polarity in the craniofacial region. Using a conditional *Wnt5a* knockout model targeting cranial neural crest-derived mesenchymal cells, we demonstrated that *Wnt5a* deficiency results in severe craniofacial defects, characterized by a reduction in the size FNP and PA1, ultimately leading to malformed distal craniofacial structures. These defects appear to be driven by disruptions in key intracellular polarity determinants, including the positioning of, primary cilia, and the Golgi apparatus, which in turn affect cell division orientation and proliferation rates. Our findings indicate that mesenchymal cells in the craniofacial region exhibit intrinsic polarity, as reflected by the spatial organization of primary cilia, centrosomes, and Golgi apparatuses along the proximal-distal (P-D) axis. This supports the notion that mesenchymal polarity mechanisms share similarities with those of epithelial cells. Mesenchymal cell polarity, however, likely involves distinct regulatory pathways because, for instance, *Vangl* genes do not seem to control Wnt/PCP pathway in the context of neural crest derivatives (Pryor et al., 2014). Loss of *Wnt5a* in our conditional knockout model disrupted this polarized character of mesenchymal cells, leading to randomization of primary cilia, centrosome, and Golgi positioning. Similar randomization of primary cilia was observed in cranial and limb chondrocytes in *Prickle1* mutants, another component of Wnt/PCP pathway (Wan and Szabo-Rogers, 2021). Proper cilia orientation is essential for accurate cell signaling and tissue formation, and misalignment of these structures can result in significant morphological defects. These findings reinforce the hypothesis that Wnt5a signaling is a critical regulator of PCP in mesenchymal cells.The disorganization of intracellular polarity components observed in *Wnt5a*-deficient mesenchymal cells likely underlies the misalignment of cell division orientation. In control embryos, the majority of mitotic spindles aligned along the P-D axis, consistent with previous studies indicating that oriented cell division plays a key role in tissue elongation and patterning (Kaucka et al., 2017; Li et al., 2017). However, in *Wnt5a* cKO embryos, cell division planes were misoriented, deviating significantly from the P-D axis. This misalignment likely contributes to the widening of the FNP and PA1 and the reduced elongation of these structures along the P-D axis.

In addition to disrupted polarity and cell division orientation, *Wnt5a* cKO mutants exhibited a significant reduction in cell proliferation, as evidenced by a lower percentage of EdU+ cells. This decrease in proliferation was particularly pronounced in the distal FNP and PA1, the regions where *Wnt5a* expression is normally strongest. The reduction in proliferation controbutes to the morphological defects observed in *Wnt5a* cKO embryos, as fewer mesenchymal progenitor cells contribute to craniofacial growth. These findings are somewhat surprising because WNT5A signaling is rather involved in PCP pathway, directional cell migration and cell differentiation. Moreover, measurement of cell proliferation rate in FNP of systemic Wnt5a KO did not indicate proliferation decrease (Kaucka et al., 2017). In several cell contexts, Wnt5a rather inhibits cell proliferation by antagonizing canonical Wnt/β-catenin signaling (Kumawat and Gosens, 2016; Mikels and Nusse, 2006; Ye et al., 2013). The molecular mechanism by which Wnt5a regulates proliferation remains unclear but may involve downstream effectors such as ROR2 and JNK signaling, which have been implicated in Wnt/PCP-mediated cell cycle regulation (Konopelski Snavely et al., 2023).

One of the key insights from our study is that mesenchymal cells in the craniofacial region exhibit a polarized organization of their Golgi apparatus and centrosomes, which is disrupted upon Wnt5a deletion. In control embryos, both organelles predominantly localized distal to the nucleus, aligning with the direction of mesenchymal cell movement and tissue elongation. This spatial arrangement is consistent with the role of Golgi and centrosome positioning in establishing front-rear polarity in migrating cells (Ravichandran et al., 2020). The shift of Golgi and centrosome positioning towards the proximal end in *Wnt5a* cKO mutants suggests that WNT5A signaling directly or indirectly regulates intracellular trafficking and cytoskeletal dynamics, processes that are crucial for directional cell migration and coordinated tissue morphogenesis. Future research should focus on identifying the specific cytoskeletal and ECM components mediating Wnt5a’s effects on mesenchymal cell polarity. Additionally, live imaging of mesenchymal cell behaviors in Wnt5a mutants could further elucidate how these defects arise dynamically during development. Understanding these mechanisms will not only enhance our knowledge of craniofacial biology but also provide potential therapeutic targets for congenital craniofacial disorders.

## MATERIALS AND METHODS

### Mouse strains

Wnt1–Cre2 (The Jackson Laboratory strain RRID:IMSR_JAX:022137) was used for specific deletion of the *Wnt5a* ^fl/fl^ gene in neural crest cells. Our standard crossing scheme for generating *Wnt5a* cKO embryos was the male Wnt1-Cre2; *Wnt5a* ^fl/+^ and the female *Wnt5a* ^fl/fl^ (RRID:IMSR_JAX:026626). Reporter line mTmG was purchased from The Jackson Laboratory (strain RRID:IMSR_JAX:007676). The presence of the vaginal plug was regarded as a embryonic day 0.5 (E0.5). Pregnant mice were anesthetized using isoflurane gas inhalation and euthanized by cervical dislocation. Embryos were dissected and placed in ice-cold PBS. All procedures involving experimental animals were approved by the Institutional Committee for Animal Care and Use (permission #PP-10/2019) with every effort to minimize the suffering and the number of animals used. Mice were genotyped as follows:

### Histology and immunohistochemistry

Embryos were harvested at specific embryonic stages and were fixed in 4% paraformaldehyde overnight at 4^0^C. After fixing with PFA, samples were prepared in two ways. After dehydration in ethanol and xylene, samples were embedded in paraffin and frontal 9-10μm sections were prepared. Alternatively, after dehydration in 30% sucrose for 24 hours, samples were embedded in OCT and 10μm or 100μm cryo-sections were obtained. For paraffin tissue sections, antigen retrieval in 0.1 M citrate buffer pH 6.0 under pressure boiling for 15 min was carried out. Tissue sections were then blocked in 5% bovine serum albumin BSA in PBS with 0.1% Triton x-100 for 2 hours and incubated overnight in primary antibody solution (1% BSA in PBS and 0.1% Triton x-100). Primary antibodies used: Sox9 (Merck Sigma AB5535), Foxf1 (R&D Systems AF4798), Msx1 (R&D Systems AF5045), Lef1 (Cell Signaling 2230), Twist1 (Santa Cruz sc-81417), Pax3 (DSHB), Tcf7L2 (Cell Signaling 2569), Arl13B (Proteintech 17711-1-AP), GM130 (Abcam ab52649), Pericentrin (Abcam ab4448). Primary antibody solutions were washed out with (PBS, Triton x-100) and incubated in fluorescent secondary antibodies solutions for one hour. Secondary antibodies: donkey anti-mouse, -rabbit, -goat with Alexa488, 568 or 647 fluorophores (Thermo Fisher Scientific). High-resolution chain reaction fluorescence in situ hybridization (HCR-FISH) was carried out according to manufacturer’s protocol (Molecular Instruments).

Immunofluorescent images were scanned by spinning disc confocal microscopy using Olympus SpinSR10 and Andor BC43 instrument, software Olympus cellSens and Fusion, respectively.

Images were processed in ImageJ-Fiji and Adobe Photoshop. Quantifications were done in Imaris software Spot module.

### Cell proliferation quantification

Pregnant mice were injected with 5-ethynyl-2’-deoxyuridine (EdU) (total 1.25 mg in 0.125 ml PBS per mouse) intraperitoneally 2 h prior to the embryo harvest. Embryos were fixed in 4% PFA in PBS overnight at 4°C, washed in PBS and dehydrated. Paraffin sections were stained with primary and secondary antibodies as described above. Sections were washed in PBT and stained using EdU BaseClick kit according to manufacturer’s instructions. Samples were scanned on Olympus SpinSR10 and Andor BC43. Quantification of double positive cells was performed in control littermates and *Wnt1-Cre2; Wnt5a* ^*fl/l*f^ embryos using Imaris software.

Percentages were calculated from the numbers of double positive cells related to the total number of DAPI^+^ cells in FNP. Values are presented as averages with standard deviations and the statistical significance was calculated using Mann-Whitney U test. P-value of <0.05 was considered significant.

### Skeletal preparations

E17.5 embryos were incubated in water for 1 hour at 4°C, scalded in boiling water for 5 minutes and they were skinned. They were dehydrated in 95% ethanol for 72 h, and the ethanol was exchanged every 12 h. Alcian blue (Sigma-Aldrich) was used to stain cartilage for 12 h, embryos were rinsed twice in fresh ethanol and incubated in ethanol overnight. Embryos were cleared in 1% KOH for 2 hours and stained with Alizarin Red S (Sigma-Aldrich) for 5 hours.

Excess of the stain was rinsed off with fresh 1% KOH for 1 h and further clearing in 1% KOH was carried out overnight. Embryos were transferred through a graded series of glycerol (25%) and KOH (1%) for 8 h, and glycerol (50%) and KOH (1%) for 48 h. Pictures were obtained using binocular microscope Olympus SYX9 and camera Olympus DP72.

## Acknowledgements

This work was supported by Czech Science Foundation (grant 22-10660S). We thank the Microscopy Service Centre of IEM CAS and Light Microscopy Core Facility, IMG CAS, Prague, Czech Republic, supported by MEYS – LM2023050 and RVO – 68378050-KAV-NPUI. This work was also funded by MEYS Czech Republic (NanoEnviCZ, LM2018124) and EU Structural Funds Pro-NanoEnviCz (CZ.02.1.01/0.0/0.0/16_013/0001821).

## Author contributions

AB designed and performed experiments and wrote manuscript. KK designed and performed experiments and wrote manuscript. LS performed experiments. OM designed and performed experiments, wrote manuscript, and provided funding. The authors declare no competing interests.

